# Novel Anammox Bacteria Discovered in the Untapped Subsurface Aquifers

**DOI:** 10.1101/2023.04.27.538623

**Authors:** Jiapeng Wu, Yuchun Yang, Ji-Dong Gu, Yiguo Hong

## Abstract

Anaerobic ammonium-oxidation (anammox) reaction is a crucial microbial nitrogen (N) transformation process contributing to the global N cycling. The currently known anammox bacteria are affiliated with two families, the marine *Ca*. Scalinduaceae and the freshwater *Ca*. Brocadiaceae, in the *Planctomycetes*. Here we report a discovery of new groups of anammox bacteria consisting of two new orders, two new families, and six new genera within the class *Candidatus* Brocadiia, all from geochemically distinct aquifer systems. Up to 25 high-quality metagenome-assembled genomes (MAGs) equipped with the core anammox metabolisms were recovered from 13 metagenomic datasets of aquifers and dominated the anammox bacterial communities in distinct aquifers unexpectedly. The comparatively smaller genome size (< 2.6 Mb) and higher coding density (> 85%) likely confer the survival advantage of them by reducing the energy consumption in cell replication and maintenance to increase the nutrient acquisition efficiency in the typically oligotrophic aquifers. Overall, the highly diversity of novel anammox bacterial lineages in aquifer systems largely expands our knowledge of the taxonomic diversity of anammox bacteria and highlights their global importance in aquifer N cycling.

---

The stoichiometry of the anammox reaction was first established in 1998 [1], in which both ammonium and nitrite are coupled to produce dinitrogen gas (N_2_) [2, 3]. To date, six candidate genera of anammox bacteria are known, including *Ca*. Scalindua [4], *Ca*. Kuenenia [5, 6], *Ca*. Brocadia [7], *Ca*. Anammoxoglobus [8], *Ca*. Jettenia [9], and *Ca*. Loosdrechtia [10]. All of them are affiliated with a monophyletic group within the order *Ca*. Brocadiales in the phylum *Planctomycetes*. Anammox bacteria are widely distributed in diverse ecosystems and play critical roles in the nitrogen (N) cycle in oxygen-depleted habitats, such as marine oxygen minimum zones [11, 12], marine sinking particles and sediments [13, 14], wastewater treatment plants [15], wetland and agricultural soils [16, 17]. Based on metagenomic sequencing, new and undescribed anammox bacteria were recently reported in subsurface ecosystems [18-20]. Subsurface aquifers, characterized by low organic carbon, nitrogen species and oxygen level but long water residence time, are ideal ecosystems for slow growing chemoautotrophic anammox bacteria [21, 22]. A growing body of evidence supports that anammox reaction plays an important role in N cycles of aquifer ecosystems [23-25]. As a result, our understanding of the phylogenetic diversity and ecological distribution of anammox bacteria remains very limited.

## Highly diverse novel anammox bacteria in aquifers

Here, we performed a systematical analysis of 13 metagenomic datasets from geochemically distinct aquifer systems, including pristine, agricultural, and seawater impacted groundwater. A total of 25 high-quality (completeness > 90%, contamination < 5%) anammox metagenome-assembled genomes (MAGs) were acquired and the full-length 16S rRNA gene sequences were recovered from 12 of these MAGs after iterative assemblies [26]. Phylogenetic analysis based on the alignment of 43 concatenated markers confirmed the placement of these 25 new MAGs into the phylum *Planctomycetes* (Figure 1a). Unexpectedly, not all of them were affiliated with the monophyletic group of the order *Ca*. Brocadiales, two new early-branching clusters were formed in the phylogenetic tree (Figure 1a). Specifically, only two MAGs (YC26 and YC27) could be assigned to the known genus *Ca*. Scalindua (Figure 1a). Eleven of them (YC19-25 and YC28-31) formed two separate clusters within the known family *Ca*. Brocadiaceae (Figure 1a, b). The full-length 16S rRNA gene sequences were recovered from MAGs YC20-23 and YC28, and the highest 16S rRNA gene identities between them and the known anammox bacteria were lower than the minimum identity cutoff value demarcating the genus level (95%) [27] (Figure 1b). Therefore, we propose clusters YC19 to YC25 and YC28 to YC31 represent two new genera within the family *Ca*. Brocadiaceae and have tentatively named them *Ca*. Wujingus (derived from Sha Seng, a figure in Chinese mythology) and *Ca*. Wunengus (derived from Zhu Bajie, a figure in Chinese mythology), respectively. Although the 16S rRNA gene identity between YC21-23 and YC28 was 95.4% (Figure 1b), slightly higher than 95%, it is believed they represent two different genera according to their apparent phylogenetical separation (Figure 1a, b). Another four MAGs (YC8-11) were assigned into the clade of the newly proposed family *Ca*. Bathyanammoxibiaceae from marine sediment [20], but phylogenetically divergent from the known genus *Ca*. Bathyanammoxibius (Figure 1a). The highest 16S rRNA gene identities between YC8 and the other anammox bacteria were lower than the minimum identity cutoff value for the genus level (95 %) (Figure 1b). Therefore, we propose the clade of YC8 to YC11 represents a new genus within the family *Ca*. Bathyanammoxibiaceae and have tentatively named them *Ca*. Avalokitesvara (derived from Guanyin Pusa, a figure in Chinese mythology). The remaining eight MAGs (YC12-17, JP1, and YC18) were phylogenetically divergent from the known families (Figure 1a, b). The full-length 16S rRNA gene sequences were acquired from YC14, YC16, JP1, and YC18 and the highest 16S rRNA gene identities between them and the known anammox bacteria were lower than the minimum identity cutoff value demarcating the order level (89 %) [27] (Figure 1b). Among them, YC18 was phylogenetically separate from cluster YC14-17 and JP1, and the 16S rRNA gene identities between them were also lower than the family cutoff 89% (Figure 1b). Therefore, we propose clusters YC14-17 and JP1 as well as YC18 represent two new orders and two new families. The order and family names of cluster YC14-17 and JP1 have tentatively named *Ca*. Hypogeohydatales and *Ca*. Hypogeohydataceae (Hypogeohydata-; derived from the Neo-Lin noun groundwater), respectively. And the order and family names of cluster YC18 have tentatively named *Ca*. Wukongales and *Ca*. Wukongaceae (Wukonga-; derived from Sun Wukong, a figure in Chinese mythology), respectively. The 16S rRNA gene identities between YC8 and cluster YC14-17 and JP1 were higher than the family cutoff 89% that the newly identified families *Ca*. Hypogeohydatales and *Ca*. Bathyanammoxibiaceae should belong to the same order *Ca*. Hypogeohydatales (Figure 1b). The 16S rRNA gene identities among YC8, JP1, YC14 & 16, and YC18 were lower than the genus cutoff 95% (Figure 1b) indicating that they represent four new genera and have tentatively been named *Ca*. Avalokitesvara (derived from Guanyin Pusa, a figure in Chinese mythology), *Ca*. Tathagata (derived from Rulai Fozu, a figure in Chinese mythology), *Ca*. Tripitaka (derived from Tang Seng, a figure in Chinese mythology), and *Ca*. Wukongus (derived from Sun Wukong, a figure in Chinese mythology), respectively. In summary, 25 high-quality anammox MAGs were acquired from distinct aquifers, which could be clustered into three orders (including two new orders), four families (including two new families), and seven genera (including six new genera) within class *Ca*. Brocadiia.

**Figure 1.**
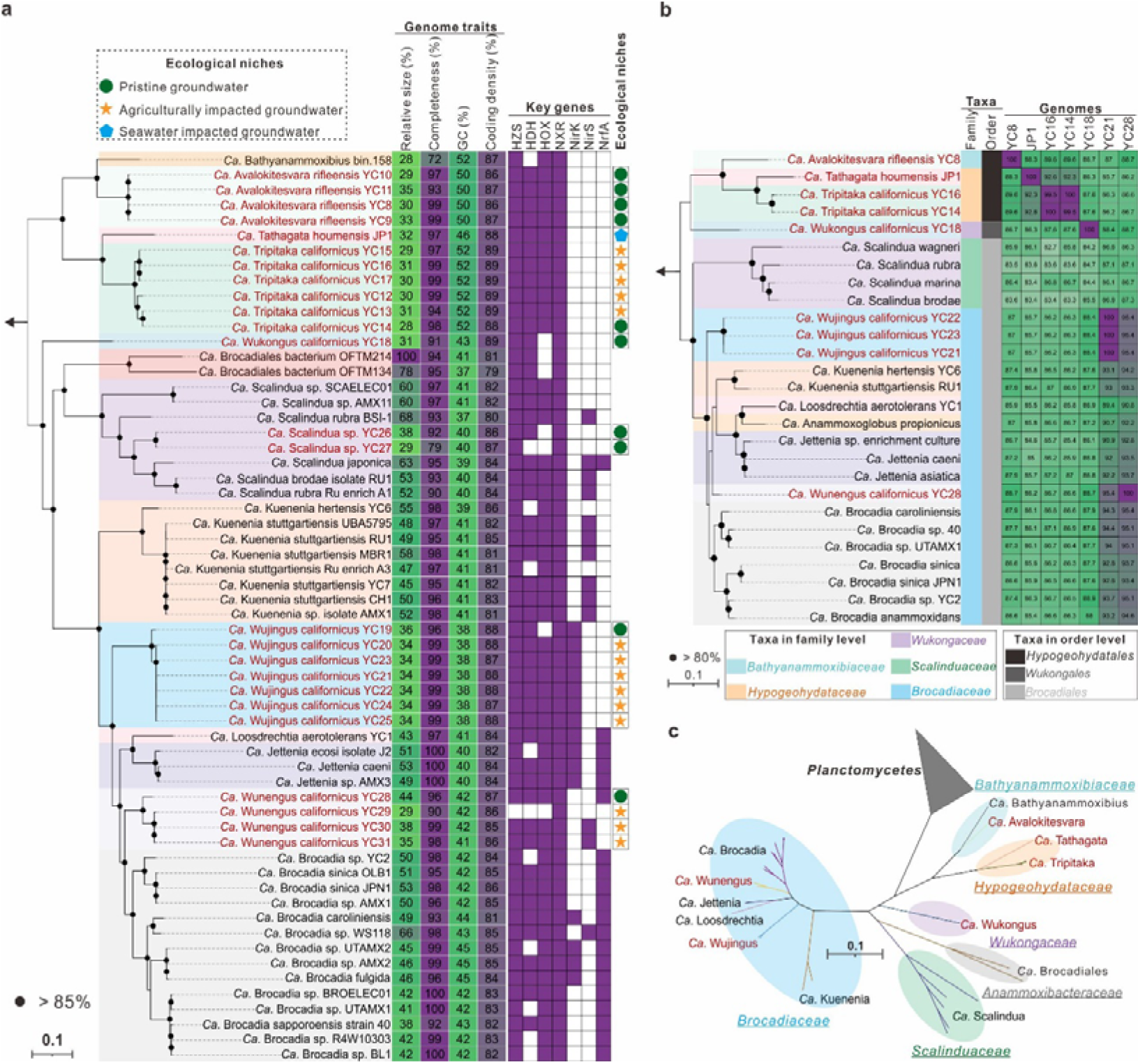
Phylogeny of the newly recovered anammox bacterial MAGs from three geochemically distinct aquifer systems, including pristine (green circles), agricultural (yellow stars), and seawater impacted groundwater (blue pentagons). (**a**) Maximum-likelihood phylogenetic tree based on the 43 concatenated markers shows the affiliations of the 25 newly recovered anammox bacterial MAGs (red font) and known anammox bacterial genomes (black font), the tree was rooted by non-anammox Planctomycetes. Bootstrap values larger than 85% are shown with a black circle. The scale bars show estimated sequence substitutions per residue. The genome traits of the novel and known anammox bacterial genomes, including relative size (%), completeness (%), GC (%) and coding density (%) are shown by heatmap. The presence and absence of key genes related to the core anammox metabolism are shown by filled and open squares, respectively. HZS, hydrazine synthase; HDH, hydrazine dehydrogenase; HOX, hydroxylamine oxidizing protein; NXR, nitrite oxidoreductase; NirK/S, NO-forming nitrite reductase; NrfA, ammonia-forming nitrite reductase. (**b**) Maximum-likelihood phylogenetic tree of the full-length 16S rRNA gene sequences recovered from the newly recovered anammox bacterial MAGs and stored in the SILVA 138 SSU database, the tree was rooted by the 16S rRNA gene sequences of non-anammox Planctomycetes. The nine representative full-length 16S rRNA gene sequences recovered from the newly acquired aquifer anammox bacterial MAGs are highlighted in red font. Bootstrap values larger than 80% are shown with a black circle. The scale bars show estimated sequence substitutions per residue. The 16S rRNA gene identities of anammox bacteria are shown by the heatmap. Anammox bacteria assigned into different families were shown by columns with different colors. (**c**) Unrooted maximum-likelihood phylogenetic trees of anammox bacteria and other non-anammox Planctomycetes based on the 43 concatenated markers. The six novel genera of anammox bacteria identified in the current study were highlighted in red font. Different families are highlighted by different background colors

## Novel genomic traits associated with niche adaptability in aquifers

Compared to the known anammox bacteria from marine and terrestrial niches, all newly acquired aquifer anammox bacteria had a relatively small genome size (2.0-3.2 Mb), especially the genomes in the new orders *Ca*. Hypogeohydatales and *Ca*. Wukongales, which were only one third of the size of the largest genome “*Ca*. Brocadiales bacterium OFTM214” (Figure 1a, Table S2). The coding density of anammox bacteria is also influenced by the ecological niche in that aquifer anammox bacteria display a significant increment in coding density (Figure 1a). The compact genome with a high coding density likely confers the survival advantage of anammox bacteria in the typical oligotrophic aquifers by reducing the energy consumption in cell replication and increasing the nutrient acquisition efficiency. Additionally, aquifer genomes in the orders *Ca*. Hypogeohydatales and *Ca*. Wukongales had a higher genomic GC content compared to genomes in the known order *Ca*. Brocadiales, even including the newly acquired *Ca*. Scalindua, *Ca*. Wujingus, and *Ca*. Wunengus MAGs from the same aquifer ecosystem (Figure 1a). It is agreed that the GC content of complex microbial communities is strongly influenced by the environment rather than phylogenetic origins [28]. But, this consensus cannot be applied to anammox bacterial community in that GC content variation is consistent well with phylogenetic diversity and varies only in a narrower range within a same genus (Figure 1a).

The biochemically unique hydrazine synthase (HZS) synthesizes hydrazine from nitric oxide and ammonium and then hydrazine oxidation is carried out by hydrazine dehydrogenase (HDH) [29-31]. Hydroxylamine, a severe inhibiter of HDH, is suggested to leak out of HZS during hydrazine production, which can be recycled back to NO by hydroxylamine oxidizing protein (HOX) [30, 32-35]. HZS, HDH, and HOX homologs related to the core anammox metabolism are highly conserved in the newly acquired aquifer MAGs, reflecting core anammox catabolism has been already equipped by aquifer anammox bacteria (Figure 2a). It is suggested that nitrite reduction (NIR) and oxidation (NXR) pathways were acquired later compared to the core anammox metabolism in the evolution[36] and distinct anammox genera utilize different NO-generating NIR [34]. The homologs of NXR were evident in all new aquifer MAGs (Figure 2a), but the putative NxrC subunits are less conserved in orders *Ca*. Hypogeohydatales and *Ca*. Wukongales, sharing 32-38% amino acid identity with the known anammox NxrC (data not shown), illustrating the significant divergence of NxrC in the evolution of anammox bacteria. Copper-containing NIR (NirK) was encoded by genus *Ca*. Wujingus but cytochrome *cd*_1_ NIR (NirS) could only be detected in some *Ca*. Wunengus MAGs (Figure 2a). However, neither NirK nor NirS coding gene could be detected in *Ca*. Avalokitesvara, *Ca*. Tathagata, *Ca*. Tripitak, and *Ca*. Wukongus, that they may harbor an unidentified enzyme to fulfill the function of nitrite reduction to NO. Pentaheme cytochrome *c* nitrite reductase (NrfAH), catalyzes the dissimilatory nitrite reduction to ammonium (DNRA) to supply anammox bacteria with ammonium, has been observed in *Ca*. Brocadia, *Ca*. Jettenia, *Ca*. Scalindua, and *Ca*. Loosdrechtia, but it was absent in the recovered aquifer MAGs with only one exception of MAG YC28 (Figure 1a). Aquifers often contain low concentrations of dissolved organic compounds [21, 22] so that anammox bacteria may not be able to outcompete heterotrophic denitrifiers and DNRA performing microbes for effective reduction of nitrite to ammonium. Therefore, it may be inferred that NrfAH coding genes have been discarded by aquifer anammox bacteria for oligotrophic niche adaptive purpose in the long evolutionary history. The absence of NrfAH indicates anammox bacteria cannot mediate DNRA to supply anammox process in the NH_4_^+^-limited aquifers. Instead, they may acquire ammonium from the coexisting microbes capable of performing DNRA and form an anammox-DNRA coupling process in the relatively nitrate-rich aquifers, which was also reported to play a major role in N loss in NH_4_^+^-limited marine systems [37].

In summary, up to 25 high-quality anammox MAGs were acquired from13 metagenomic datasets of distinct aquifers in this study, which could be clustered into three orders (including two new orders), four families (including two new families), and seven genera (including six new genera) within class *Ca*. Brocadiia. The relatively small genome size and high coding density likely confer a selective advantage to aquifer anammox bacteria in the typical oligotrophic aquifer systems. The highly diverse of novel anammox bacteria identified in aquifers with quite different genomic traits substantially expand our understanding on the phylogenetic diversity of anammox bacteria. Future enrichment and investigation of the key metabolisms and survival strategies will lead to a better understanding of the niche adaptability and ecological function of these novel anammox organisms, with implications to the potential new biotechnology of groundwater and wastewater treatments.

## Materials and Methods

### Sampling collection and characterization

Sediment and groundwater samples were collected from a coastal aquifer (115.15°E, 22.81°N) adjacent to the South China Sea, located in Houmen Town, Shanwei City (Table S1). One sediment core (80 cm in length) was sampled at the high tide on 3 December 2019 by using a Portable soil core sampler (Rhino S1, USA) and divided at 10 cm intervals over the vertical depth. Each section of the sediments was immediately sealed in plastic bags, placed in a cooler box and transported back to the laboratory. Sediment samples in the depth of 60 cm (near the water table) were selected for DNA extraction and metagenomic sequencing. DNA was extracted from 20 g moist sediment using PowerSoil DNA Isolation Kit (Qiagen, USA) and then sequenced using Illumina Novaseq 6000. Groundwater was collected corresponding to the core sample depths by peristaltic pumps (GeoTech, USA) and groundwater collection units (GVPKIT, ASM, USA). Groundwater properties including dissolved oxygen, salinity, and dissolved nitrogen contents were also measured followed the method reported elsewhere [38].

### Nucleic acid extraction, sequencing, and genome binning

Seawater impacted groundwater sample was collected from Houmen, China and DNA was extracted for metagenomic sequencing (100 Gb data). Metagenomic datasets of pristine and agricultural impacted groundwater in Rifle [39] and Modesto [40], United States were downloaded from NCBI (BioProject: PRJNA268031, PRJNA288027, and PRJNA640378). DNA extraction, sequencing, de novo metagenomic assembly, and binning were previously described [15]. In total, 46 putative aquifer anammox genomes were recovered after assembly and binning based on the taxonomic assignments of GTDB-Tk [41]. The qualities of these genomes were further improved through an iterative assembly as described in a previous study [26].

### Genome annotation and phylogenetic analyses

CheckM v1.0.6 [42] was used to produce an alignment of 43 concatenated markers from the 25 newly acquired high-quality anammox genomes, 33 known anammox genomes, and other 157 genomes from the PVC superfamily. 16S rRNA genes of anammox bacteria stored in the SILVA 138 SSU database were used for phylogenetic analysis with the full-length or nearly full-length 16S rRNA genes in the 25 anammox genomes. HzsA sequences from previously published anammox bacteria and the 25 newly acquired anammox genomes were used for phylogenetic analysis. All above sequences used for phylogenetic analysis were aligned by MAFFT online service [43], and gaps in the multiple sequence alignment were removed by trimAl V1.2 with the setting ‘-gt 0.1’. Maximum likelihood trees were built using IQ-TREE with the default settings[44] and visualized using iTOL [45]. The identities of full-length 16S rRNA gene sequences of anammox bacteria were calculated by Blast 2 [46].

## Data availability

Data of this study are available from the authors.

### Acknowledgement

We thank Yiben Li, Xiaohan Liu and Prof. Bongkeun Song for their assistance with field work; Jiaqi Ye and Cuihong Jiang for their assistance with data analysis. We thank the help from Dr. Rongfeng Cui in Latinized naming. This study was funded by the National Natural Science Foundation of China (No. 42006122, No. 91851111, No. 32100086, No. 92051103, and No. 31870100), Basic and Applied Basic Research Foundation of Guangdong Province (No. 2020A1515110597, No. 2019B1515120066, No. 2020A1515111033, No. 2021A1515011195), Basic and Applied Basic Research Foundation of Guangzhou City (202102021232) and Young Talent Research Project of Guangzhou Education Bureau (No. 202032795).

## Author contributions

J.W., and Y.H. designed the study and collected the samples. J.W. extracted DNA and performed assembly and binning. Y.Y. and J.W. performed bin selection, bin reassembly, phylogenetic analysis, abundance and community composition analysis, and MAGs annotation. J.W., Y.Y. drew the figures and drafted the manuscript. Y.H. and J.G. revised and edited the manuscript. All authors reviewed the results and approved the manuscript.

## Competing interesting

The authors declare no competing interests.

